# Diversity analysis of soil microbial population abundance before and after planting JunCao “Oasis No. 1” in saline-alkali soil

**DOI:** 10.1101/2021.10.26.466034

**Authors:** Xiao Zhi Qi, Lin Zhan Xi

**Affiliations:** National JunCao engineering technology center of Fujian agriculture and Forestry University, Fujian, Fuzhou, China

**Keywords:** JunCao “Oasis No.1”, Rhizosphere soil, 16sDNA high-throughput sequencing, α -diversity analysis, β-diversity analysis, OTU analysis, PcoA analysis, PCA analysis, NMDS analysis

## Abstract

In order to explore the difference of soil microbial population structure and abundance before and after planting JunCao “Oasis No. 1” in saline-alkali soil, verify the improvement effect of JunCao “Oasis No. 1” on microbial population structure and abundance in saline-alkali soil. Samples were collected from the blank saline area with and without JunCao “Oasis NO.1” and no plant growth on the surface, respectively, as Experimental group soil samples (S.Y.1-S.Y.8) and Blank group soil samples (K.B.1-K.B.8).16sDNA high-throughput sequencing technology was used for sequencing analysis respectively, and the diversity of microbial population abundance between them was compared and analyzed.The results showed that the diversity of microbial population abundance in the experimental group was significantly higher than that in the blank group, and the diversity of microbial population abundance in the experimental group was significantly different from that in the blank group, indicating that the composition of microbial population in the experimental group was significantly different from that in the blank group. In the OTU cluster analysis, the number of OTU clusters in the Experimental group soil samples (S.Y.1-S.Y.8) was significantly higher than that in the Blank group soil samples (K.B.1-K.B.8). In the sample complexity analysis of α-diversity analysis, the richness and diversity of microbial population in soil samples of Experimental group (S.Y.1-S.Y.8) were significantly higher than that in soil samples of Blank group (K.B.1-K.B.8), which was clearly reflected in the Species accumulation boxplot and Graph of species diversity. In the β-diversity analysis, PcoA, PCA and NMDS analysis methods were used to analyze the difference of microbial population diversity between Experimental soil samples (S.Y.1-S.Y.8) and Blank soil samples (K.B.1-K.B.8). The results showed that the diversity of microbial population in Experimental soil sample (S.Y.1-S.Y.8) was significantly different from that in Blank soil sample (K.B.1-K.B.8). In this paper, 16sDNA high-throughput sequencing technology was used to analyze the diversity of microbial population abundance between Blank soil samples and Experimental soil samples, and it was proved that JunCao “Oasis No. 1” had good saline-alkali soil improvement characteristics. It can effectively increase the abundance of microbial population in saline-alkali soil, so as to restore the microbial population ecosystem in saline-alkali soil, which has important application value in soil saline-alkali control.

---

Soil salinization refers to the constant accumulation of salt ions in soil, which leads to increasing soil alkalinity^[1]^. It is a major ecological problem facing human society, which has seriously affected the survival and development of human beings^[2]^.

The most significant feature of saline-alkali soil is the increase of soil alkalinity, which can lead to the precipitation of soil salts from the inside of the soil and cover the surface of the soil, resulting in the appearance of frosty white soil. At the same time, soil compaction is also accompanied by soil particles condensed into blocks due to the interaction between soil ions, resulting in soil compaction^[3]^. Soil compaction can seriously damage the soil structure, resulting in the change of soil properties^[4]^.

The most serious problem of soil in saline-alkali land is not only soil compaction, which leads to the complete change of soil properties, but also the complete destruction of soil microbial community ecosystem^[5]^. After the microbial population ecosystem in saline-alkali soil was destroyed, the microbial metabolism and other life activities were greatly reduced, which led to the loss of the internal restoration mechanism of soil fertility in saline-alkali soil^[6]^.

Therefore, restoring the microbial community ecosystem in saline-alkali soil has become the core issue of soil remediation and management in saline-alkali soil^[7]^. At present, the soil restoration and treatment methods of saline-alkali land are mainly physical and chemical methods. The physical method is mainly to wash the soil of saline-alkali land by digging ditches, so as to filter the alkaline salt ions in the soil of saline-alkali land. Chemical method is mainly through adding various soil neutralizer to the soil of saline-alkali land, adsorb and neutralize the alkaline salt ions in the soil of saline-alkali land, so as to reduce the alkalinity of the soil of saline-alkali land.

However, both physical and chemical methods have serious side effects when treating saline-alkali soil, which will cause secondary damage to soil^[8]^. Physical excavation of ditches increases the destruction of the soil’s internal structure and the loss of organic matter and other soil nutrients. Chemical methods in the addition of soil neutralizer at the same time, will cause neutralizer and soil alkaline ion reaction generated sediment in the soil deposition, thereby destroying the original soil ion distribution in the soil, aggravating soil compaction, and even lead to serious secondary problems intensified by soil heavy metal ion deposition. Therefore, using biological methods to select new plants with good soil improvement characteristics of saline-alkali soil for biological treatment of saline-alkali soil has become the primary choice for future saline-alkali soil treatment^[9]^.

As a new plant cultivated by Professor Lin Zhanxi of Fujian Agriculture and Forestry University in 1987, JunCao was first used to replace trees to cultivate various kinds of edible and medicinal fungi, so as to alleviate the contradiction of mycelium forest caused by large area cultivation of edible and medicinal fungi. After more than 20 years of development, JunCao has been widely used in the field of environmental ecological management, among which the improvement and management of saline-alkali land is the most significant. JunCao “Oasis No. 1” is a new type of grass specially bred for the treatment of saline-alkali soil, which has achieved remarkable treatment effect.

Practice has proved that the JunCao “Oasis No. 1” can effectively improve the abundance of microbial population in saline-alkali soil, restore the microbial population ecosystem in saline-alkali soil, enhance microbial metabolism and other life activities in saline-alkali soil, and thus effectively restore soil fertility in saline-alkali soil. It has good soil management ability of saline-alkali land.

In this paper, 16sDNA high-throughput sequencing technology was used to analyze the microbial population of the rhizosphere soil of JunCao “Oasis No.1” and the blank lot of JunCao “Oasis No.1”, combined with α-diversity analysis, β-diversity analysis, OTU analysis, PcoA analysis, PCA analysis and NMDS analysis. By analyzing the difference of microbial population abundance between the two species, it is proved that the JunCao “Oasis No.1” can effectively improve the microbial population abundance in saline-alkali soil, and has good soil governance ability and important ecological application value.

## 1 Materials and Methods

### 1.1 Collection of soil samples

The collection of soil samples was divided into Blank group and Experimental group. The Blank group was saline-alkali soil in blank area without planting JunCao “Oasis No. 1”, and the Experimental group was rhizosphere soil of JunCao “Oasis No. 1”. The five-point sampling method was used to collect soil samples. The sampling depth of both soil samples was 20cm, and 8 soil samples were collected in each group. Soil samples in Blank group were numbered K.B.1-K.B.8, and soil samples in Experimental group were numbered S.Y.1-S.Y.8 (Table 1).

**Table 1.** Soil sample collection information.

| The sample<br>Group | Sampling<br>Method | Sampling<br>Depth | Sample<br>Location | Sample<br>Number |
| --- | --- | --- | --- | --- |
| Blank soil<br>sample | Five point<br>sampling method | 20 cm | Blank saline<br>area | K.B.1-K.B.8 |
| Experimental soil<br>sample | Five point<br>sampling method | 20 cm | Rhizosphere of<br>Juncao | S.Y.1-S.Y.8 |

### 1.2 Extraction of total microbial DNA from soil samples

According to the operating instructions of MP Biomedicals Fast DNA soil sample extraction kit, total microbial DNA of Experimental group soil sample and Blank group soil sample is extracted. After PCR amplification, the product is purified and the DNA library is prepared, and the DNA library is sequenced by machine after passing the quality test.

### 1.3 16SrDNA high-throughput sequencing analysis

According to the characteristics of the amplified 16S region, a small fragment library was constructed and double-terminal sequencing was performed on Illumina NovaSeq sequencing platform^[10]^. Using Reads splicing filter and OTUs (Operational Taxonomic Units) clustering, species annotation and abundance analysis were performed for soil-like microbial populations in the Experimental and Blank groups. Through Alpha-Diversity and Beta-Diversity analysis, the differences in microbial population abundance between the Experimental group and the Blank group were revealed for personalized analysis and in-depth data mining.

## 2 Results and analysis

### 2.1 OTU analysis

In order to study the species composition of each sample, Effective Tags of all samples were clustered for OTUs (Operational Taxonomic Units) with 97% Identity, and then species annotation was made for the sequences of OTUs.

#### 2.1.2 Venn Graph analysis based on OUT

According to the OTUs results obtained by clustering and research requirements, the common and unique OTUs among different samples of the Experimental group and the Blank group were analyzed. When the sample number is greater than 5, a Venn Graph is drawn.

According to the Venn diagram of OUT between different samples of soil samples in the Experimental group and Blank group (FIG. 1), the number of OUT sequences in all samples in the Experimental group was significantly higher than that in the Blank group, and there were common OUT sequences between all samples in the Experimental group and Blank group.

**FIG. 1.**
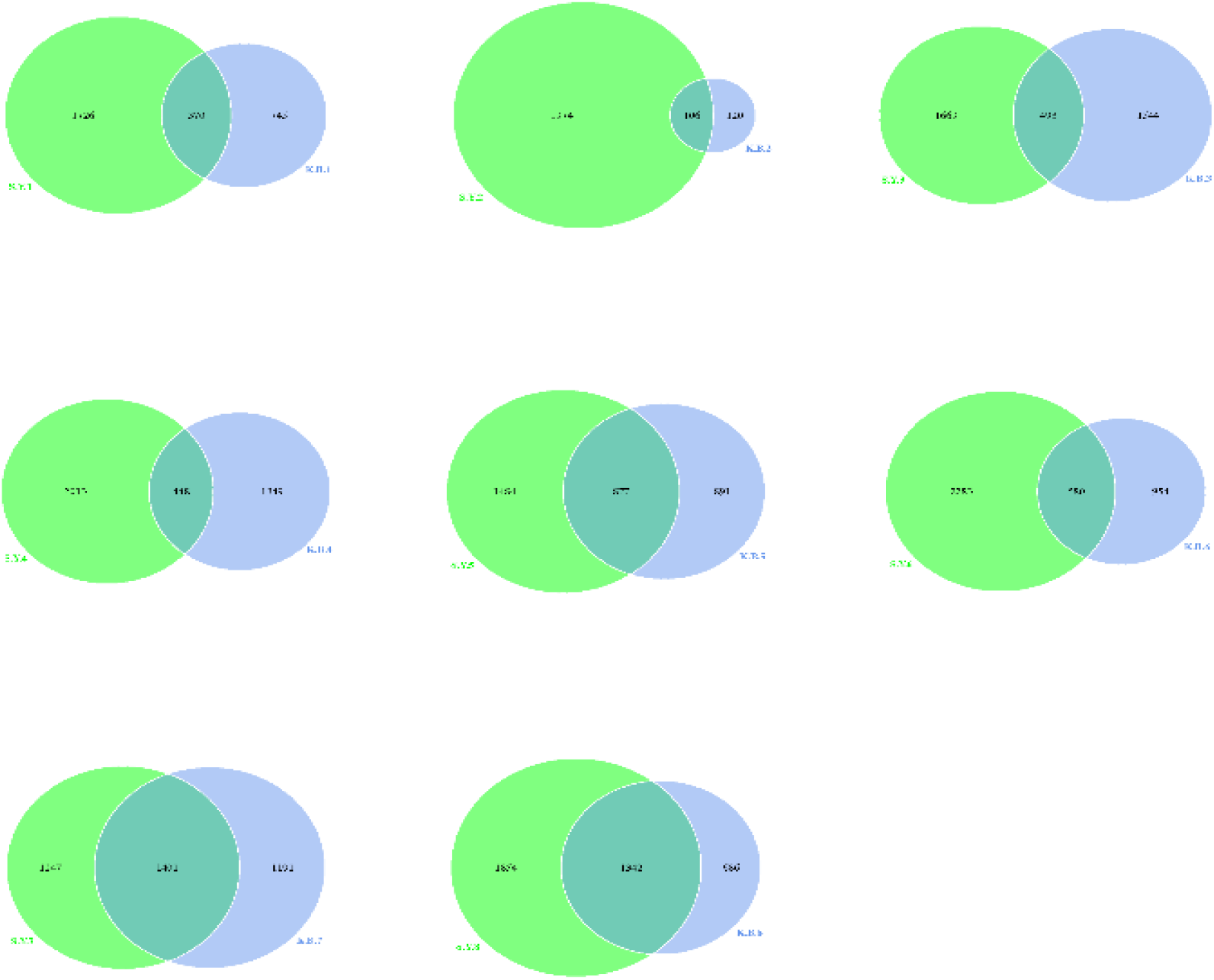
Venn graph of different samples

Among them, the samples with the largest difference in the number of OUT sequences were S.Y.2 samples in the Experimental group and K.B.2 samples in the Blank group. The number of OUT sequences of S.Y.2 sample was 1974, the number of OUT sequences of K.B.2 sample was 120, the number of OUT sequences difference was 1854, and the number of OUT sequences shared by the two samples was 106.

The samples with the smallest difference in the number of OUT sequences were S.Y.7 in the Experimental group and K.B.7 in the Blank group. The number of OUT sequences in S.Y.7 and K.B.7 samples was 1247, 1191 and 56, respectively. The number of OUT sequences in both samples was 1401.

#### 2.1.3 Phyloevolutionary relationship analysis

In order to further study the phylogenetic relationships of microbial populations at the genus level of each sample in the Experimental group and the Blank group, the phylogenetic relationships of representative sequences of Top100 genera were obtained through multi-sequence alignment to display the phylogenetic relationships of these genera. The colors of branches in the developmental tree indicate their corresponding phylum, and each color represents a phylum

According to the phylogenetic tree of each sample’s genus level population (FIG. 2), it can be concluded that the dominant phyla developed in each soil sample at the genus level are Proteobacteria and Firmicutes, and Proteobacteria is the dominant phylum. Proteobacteria was the second dominant phylum. The two dominant phyla developed into two large branching groups.

**FIG. 2.**
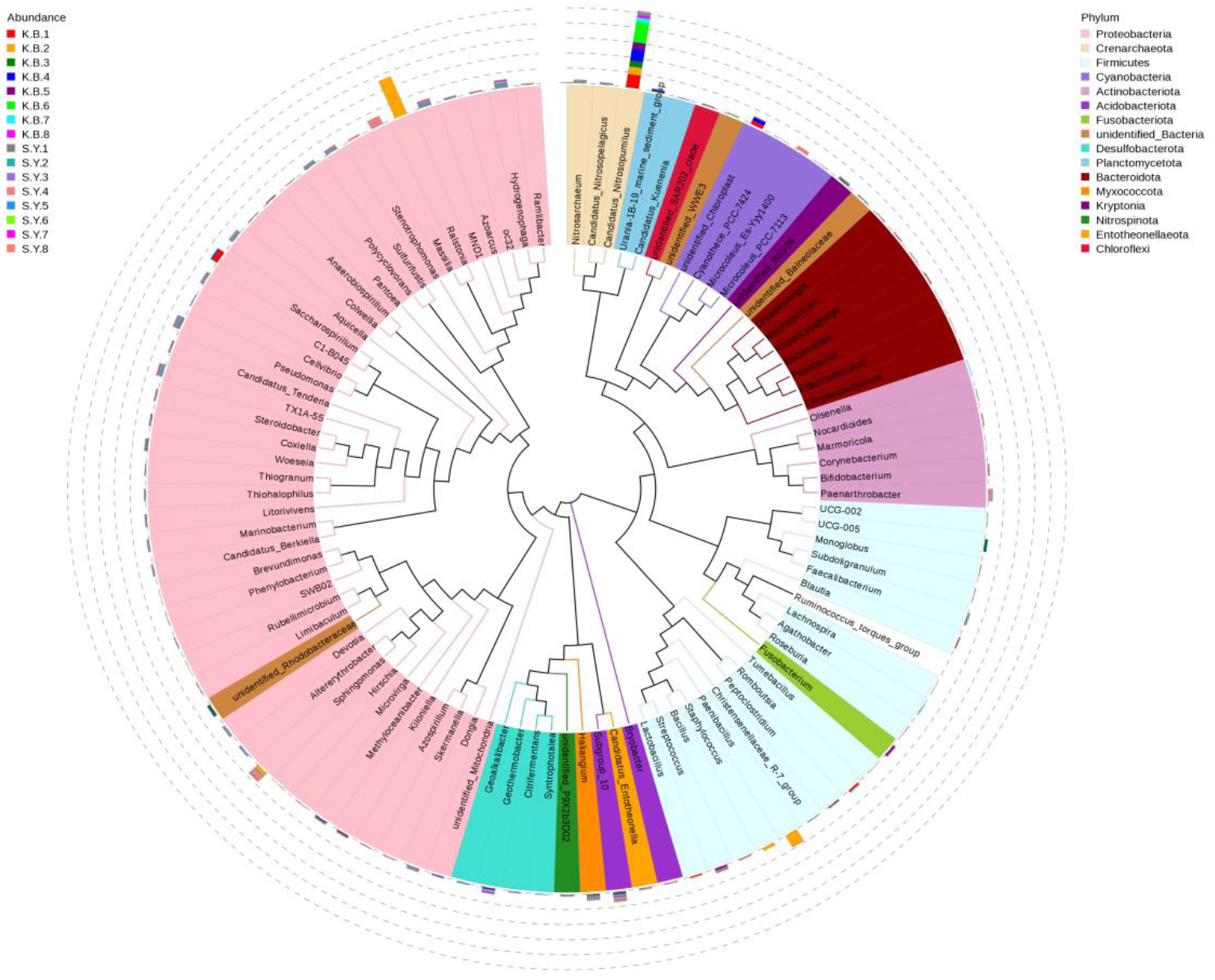
Phylogenetic tree of genus level population

Among Proteobacteria, Pseudomonas is the dominant genus in the soil samples of the Experimental group, and there is a large number of Pseudomonas in the Experimental group. The dominant position of Sphingomonas was not obvious, and only S.Y.4 sample from the experimental group had a large distribution in Sphingomonas.

Among the Firmicutes, Bacillus was the dominant bacterium in the soil samples of the Experimental group, and there was a large number of Bacillus in the soil samples of the Experimental group. The soil samples in the Experimental group were abundant in Streptococcus. At the same time, the K.B.2 samples in blank group soil samples were widely distributed in Staphylococcus and Paenibacillus.

### 2.2 α-Diversity analysis

Alpha-Diversity analysis, also known as sample complexity analysis, is used to analyze the Diversity of microbial community Within the sample^[11]^. A single-sample diversity analysis (Alpha diversity) can reflect the richness and diversity of microbial communities within a sample, including the use of a Species Accumulation Boxplot, Species Diversity Curve and a series of statistical analysis index to assess the differences in species richness and diversity of microbial communities within a sample.

#### 2.2.1 Alpha-diversity index analysis

The Alpha-Diversity analysis index (Shannon, Simpson, Chao1, ACE, Goods_coverage, PD_whole_tree) of soil samples in each Experimental group and Blank group under 97% consistency threshold was statistically analyzed. Generate Alpha Indices (Table. 2), the amount of data selected during homogenization: cutoff=28950.

**Table. 2.**
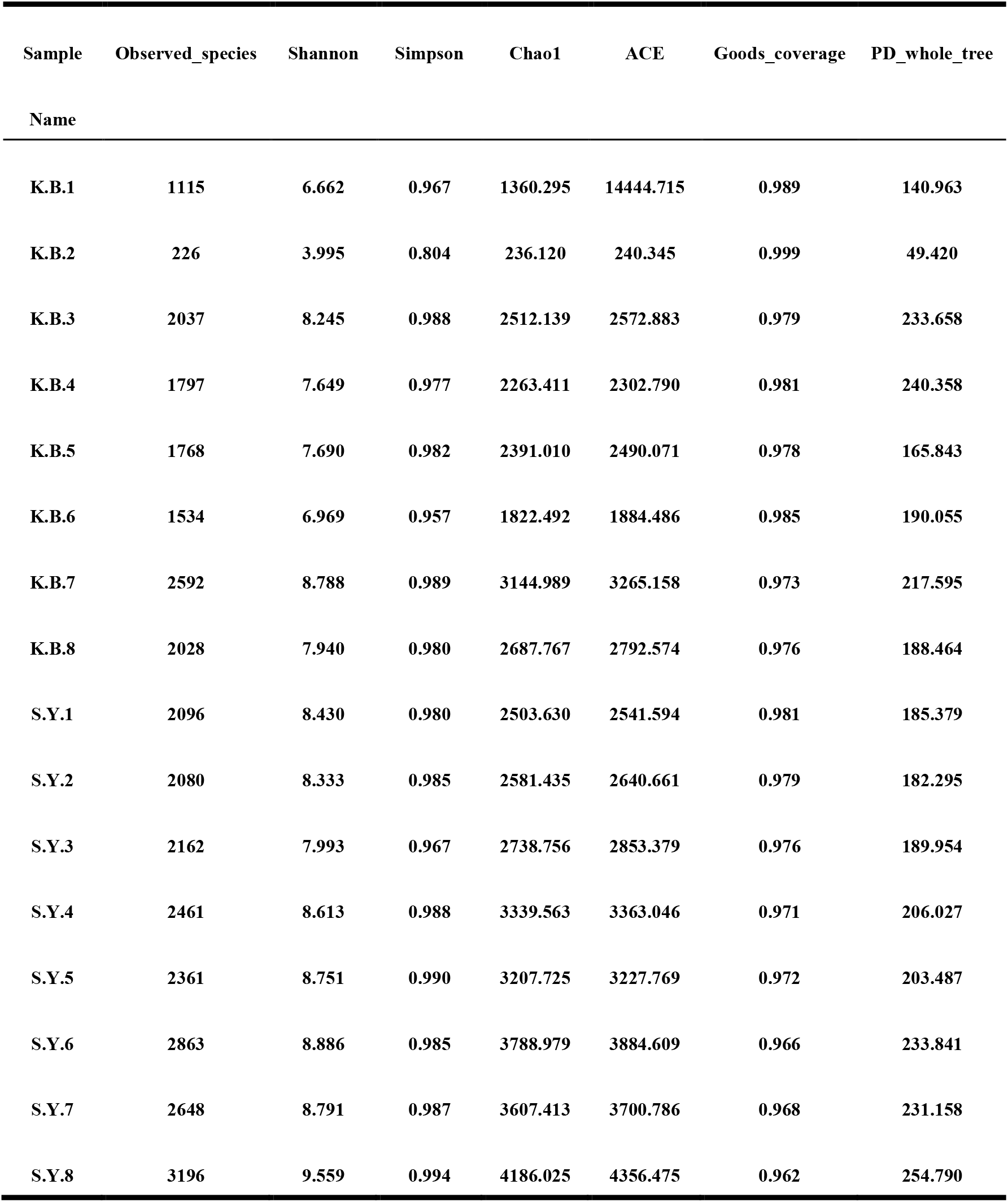
Alpha Indices.

Where Observed_species represents the intuitively observed number of species (OTUs number). Shannon represents the total number and proportion of classification in the sample. The higher the community diversity was, the more uniform the species distribution was and the larger Shannon index was. Simpson is used to characterize the diversity and evenness of species distribution in the community. Chao1 are the total number of species included in the estimated community samples. ACE is the estimated number of OTU in the community. Goods_coverage stands for sequencing depth index. PD_whole_tree represents the kinship of species in the community. Simpson has three display forms, namely, Simpson’s Index (D), Simpson’s Index of Diversity (1-D) and Simpson’s Reciprocal Index (1 / D), all of which are used to reflect community Diversity. The effect is similar but the result of the calculation is different. This analysis uses Simpson’s Index of Diversity (1-D).

Table. 2 shows that the Observed_species value, Shannon index, Chao1 value and ACE value of samples in the Experimental group are higher than those in the Blank group, indicating that the microbial population abundance in the Experimental group is higher than that in the Blank group.Meanwhile, the PD_whole_tree value of all samples in the Experimental group was higher than that in the Blank group, indicating that the genetic correlation between microbial species in the Experimental group was higher than that in the Blank group. There was no significant difference between Simpson values of samples in the Experimental group and samples in the Blank group, indicating that the composition of microbial populations in the Experimental and Blank groups were stable and continuous.

#### 2.2.2 Analysis of Species Diversity Curve

Rarefaction Curve and Rank Abundance are common curves describing the diversity of samples within groups. The Rarefaction curve is to randomly extract a certain amount of sequencing data from the sample, count the number of species they represent (OTUs number), and build the curve with the amount of sequencing data extracted and the number of corresponding species. Rarefaction curve can directly reflect the rationality of the sequencing data amount and indirectly reflect the richness of species in the samples. When the curve tends to be flat, it indicates that the sequencing data amount is progressive and reasonable, and more data amount will only generate a few new species (OTUs).

The Rank Abundance curve is to sort OTUs in the sample from large to small according to the relative abundance (or the number of sequences contained) to get the corresponding sequence number. Then, the ordinate is the ordinate of the sequence number of OTUs and the ordinate is the relative abundance of OTUs (or the relative percentage of sequence number in the OTU of this grade), and these points are connected by broken lines. That is, Rank Abundance curve can be drawn, which can intuitively reflect the richness and evenness of species in samples. In the horizontal direction, species richness was reflected by the width of the curve. The higher the species richness, the larger the span of the curve along the horizontal axis. In the vertical direction, the smoothness of the curve reflects the uniformity of species in the sample. The flatter the curve, the more uniform the species distribution^[12]^.

In the Rarefaction Curve, the horizontal coordinate is the number of sequencing lines randomly selected from a sample, and the vertical coordinate is the number of OTUs that can be constructed based on the number of sequencing lines to reflect the sequencing depth. Different samples are represented by curves in different colors. In the Rank Abundance curve, the abscissa is the serial number sorted by OTUs abundance, and the ordinate is the relative abundance of corresponding OTUs. Different samples are represented by broken lines in different colors.

According to the Rarefaction Curve of microbial species of each soil sample (FIG. 3), the number of OTU of microbial species in each sample of the Experimental group was significantly higher than that of the samples of the Blank group. The highest number of OTU of microbial species in Experimental soil samples was S.Y.8, and the lowest number was S.Y.2. K.B.2 was the sample with the lowest OTU number of microbial species in the Blank group, and the OTU number of microbial species was much lower than other samples. The highest sample was K.B.3, but the number of OTU of microbial species in K.B.3 was still lower than that in S.Y.2.

**FIG. 3.**
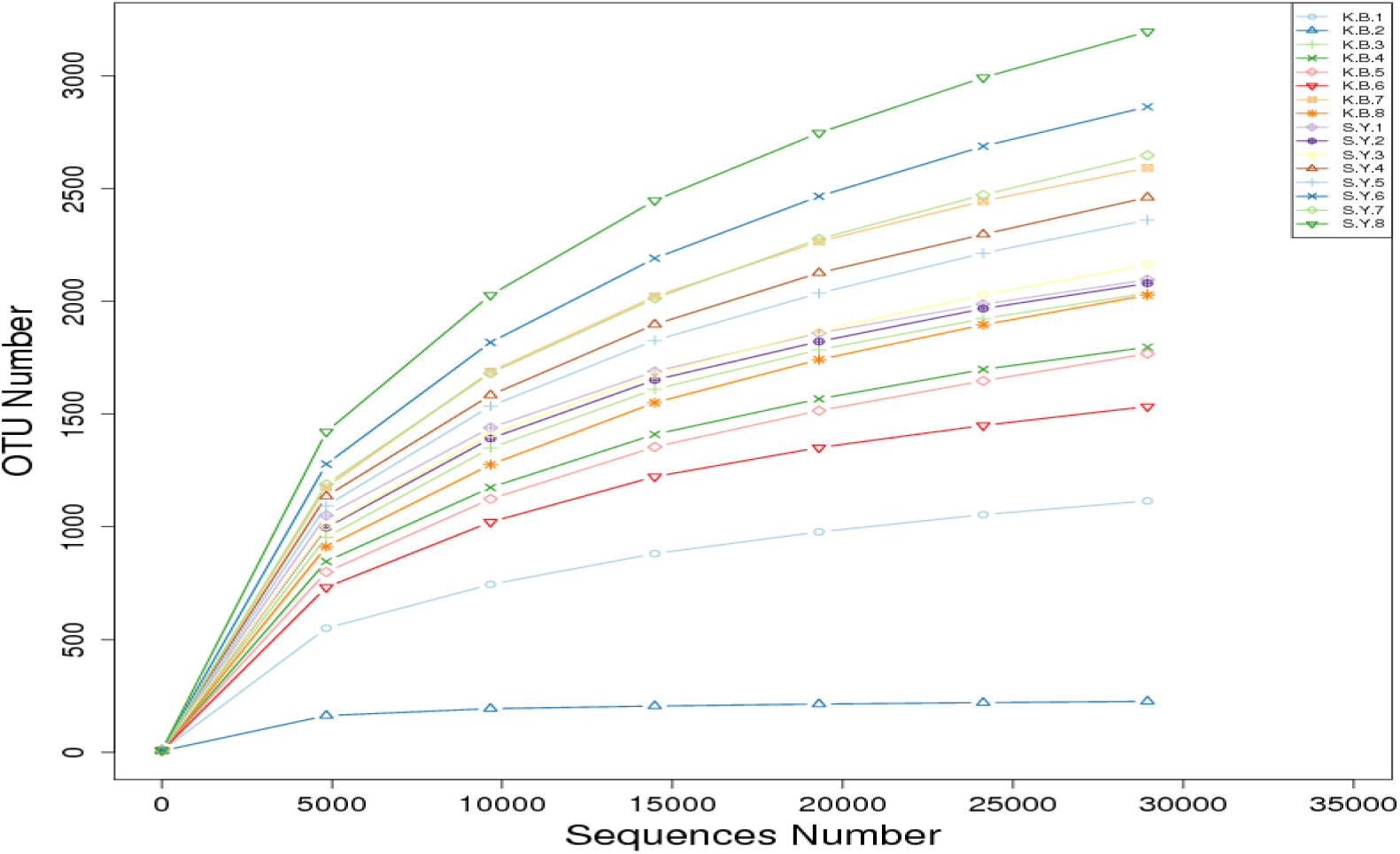
Species Rarefaction Curve

This indicates that the abundance of microbial population in the Experimental group samples is much higher than that in the Blank group samples, and the number of microbial species in the Experimental group soil samples has been effectively restored and improved.

According to the Rank Abundance curves of microbial species of each soil sample (FIG. 4), the span of each sample in the Experimental group on the horizontal axis was significantly higher than that of each sample in the Blank group. In the Experimental group, the sample with the largest span on the horizontal axis is S.Y.8, and the sample with the smallest span is S.Y.3. In the Blank group, the sample with the largest span along the horizontal axis is K.B.3, but the span is still smaller than that of S.Y.3. The sample with the smallest span on the horizontal axis is the K.B.2 sample, which has a much smaller span than the other samples.

**FIG. 4.**
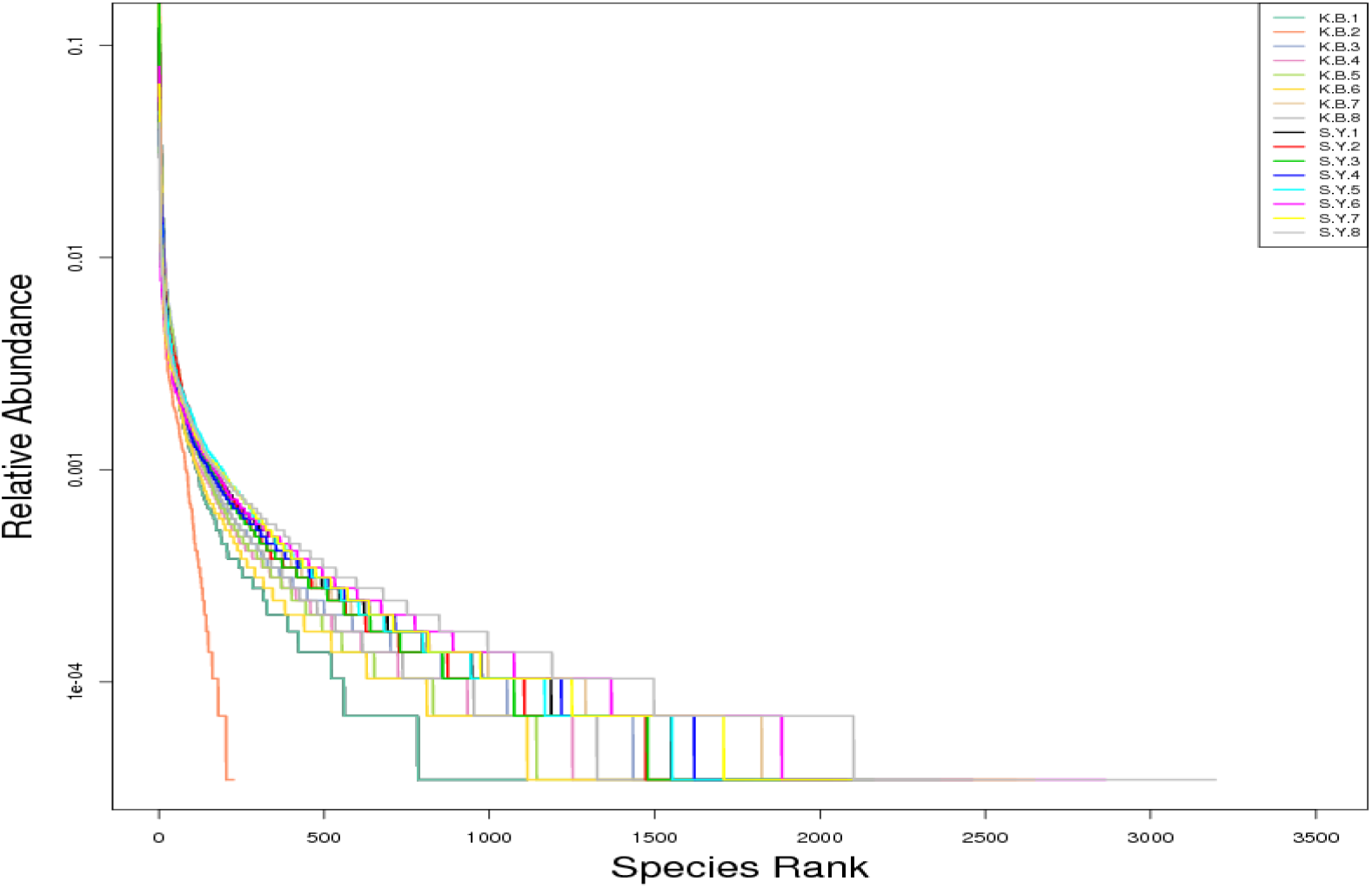
Rank Abundance curves of microbial species

In the vertical direction, the smoothness of each sample in the Experimental group was significantly higher than that in the Blank group. In the Experimental group, the sample with the most gentle decline is S.Y.8, and the sample with the least gentle decline is S.Y.3. In the Blank group, the sample with the most gentle decline was K.B.3, but the gentle decline was still smaller than that of S.Y.3. In the Blank group, the sample with the smallest decline gentleness is K.B.2, which is much smaller than other samples.

This indicates that the microbial species richness of samples in the Experimental group is significantly higher than that in the Blank group, and the uniformity of microbial population species composition in the Experimental group is significantly higher than that in the Blank group, and the microbial population species composition in the Experimental group is more uniform and stable than that in the Blank group.

#### 2.2.3 Species Accumulation Boxplot analysis

Species Accumulation Boxplot is an analysis describing the increase of Species diversity with the increase of sample size. It is an effective tool for investigating Species composition and predicting Species abundance in samples. It is widely used to judge the adequacy of sample size and estimate species richness.

The abscissa of the Species Accumulation Boxplot is the sample size. The y-coordinate is the number of OTU after sampling. The results reflect the rate at which new OTUs (new species) emerge under continuous sampling. In a certain range, with the increase of sample size, if the box plot position showed a sharp rise, it indicated that a large number of species were found in the community. When the box plot position tends to flat, it indicates that the species in this environment will not increase significantly with the increase of sample size.

According to the Species Accumulation Boxplot of each sample, the growth rate of new species began to increase with the increase of sample number, and the number of new species in each soil sample in the Experimental group was significantly higher than that in the Blank group. As the sample size increases further, the rate of new species growth slows down and tends to saturation.

This indicates that the microbial species richness of samples in the Experimental group is significantly higher than that in the Blank group. The microbial population in the Experimental group soil sample is more complex than that in the Blank group, and contains more new microbial populations.

### 2.3 β-Diversity analysis

β-Diversity analysis, also known as Beta Diversity, is a comparative analysis of the composition of microbial communities in different samples. Firstly, according to the species annotation results and OTUs abundance information of all samples, the OTUs information of the same classification was merged to obtain the Profiling Table. Meanwhile, the systematic relationship between OTUs is used to further calculate the Unweighted Unifrac^[13,14]^.

Unifrac distance is an inter-sample distance calculated by using the evolutionary information between microbial sequences in each sample. If you take more than two samples, you get a distance matrix. Then, the Weighted Unifrac distance was further constructed by using OTUs abundance information for Unweighted Unifrac^[15]^. Finally, multivariate statistical methods such as Principal Component Analysis (PCA), Principal Coordinate Analysis (PCoA), Non-Metric Multi-dimensional Scaling (NMDS), Unweighted Pair-group Method with Arithmetic Means analysis (UPGMA) and β-Diversity index difference analysis were used to find the differences among different samples.

#### 2.3.1 Distance matrix heatmap

In the β-Diversity study, Weighted Unifrac distance and Unweighted Unifrac distance were used to draw a Heatmap to measure the dissimilarity coefficient between the two samples. The smaller the value, the smaller the difference in species diversity between the two samples.

Weighted Unifrac distance and Unweighted Unifrac distance were used to draw the Heatmap, and the numbers in the squares were the dissimilarity coefficients between two samples. The smaller the dissimilarity coefficient was, the smaller the difference of species diversity was. In the same grid, the upper and lower values represent Weighted Unifrac and Unweighted Unifrac distance respectively.

According to the heatmap of β-Diversity index of each sample, there were significant differences in microbial species diversity between each sample of the Experimental group and the Blank group. The difference in species diversity between S.Y.3 in the Experimental group and K.B.2 in the Blank group was the greatest.

This indicates that the microbial population composition of the Experimental group samples has been significantly different from that of the Blank group samples. The abundance of microbial population of the Experimental group soil samples is significantly higher than that of the Blank group soil samples, and there are new microbial populations different from that of the Blank group soil samples.

#### 2.3.2 Dimension reduction analysis

##### 2.3.2.1 PCoA analysis

PcoA Analysis, also known as Principal Co-ordinates Analysis^[16]^, is an Analysis method that extracts the most important elements and structures from multidimensional data through a series of feature values and feature vector sorting. This method conducts PCoA analysis based on Weighted Unifrac distance and Unweighted Unifrac distance, and selects the principal coordinate combination with the largest contribution rate for diagram display. The closer the sample distance was, the more similar the species composition structure was. Therefore, samples with high similarity in community structure tended to cluster together, while samples with great community difference were far apart.

The PCoA analysis diagram can be presented in two forms: two-dimensional and three-dimensional. The two-dimensional PCoA diagram selects the first and second principal coordinates for display. While the Three-Dimensional PCoA diagram selects three main coordinates to display, and the coordinates can be flexibly adjusted. This analysis adopts the two-dimensional display form. The abscissa in the PcoA analysis diagram represents one principal component, the ordinate represents another principal component, the percentage represents the contribution value of the principal component to the sample difference, each point in the diagram represents a sample, and the samples of the same group are represented by the same color.

According to the PCoA analysis diagram based on Weighted Unifrac distance, the composition of microbial populations of each sample in the Experimental group and the Blank group showed obvious differentiation. With the exception of S.Y.4 sample (which may be caused by uneven distribution of soil microbial population), all other samples in the Experimental group showed similarity and concentrated distribution of loci.

According to the PCoA analysis diagram based on Unweighted Unifrac dis tance, the composition of microbial populations of samples in the Experimental group and Blank group showed obvious differences. All samples in the experi mental group showed a high degree of similarity, and the distribution of loci was very dense.

The loci distribution of S.Y.1, S.Y.2, S.Y.3, S.Y.4 and S.Y.5 samples were very concentrated and close to each other, showing a high degree of similarity. The loci of S.Y.6 and S.Y.7 samples were closely distributed, showing a high degree of similarity.

##### 2.3.2.2 PCA analysis

PCA Analysis is also called Principal Component Analysis^[17]^. It is a method of applying variance decomposition based on Euclidean distances to reduce dimension of multidimensional data so as to extract the most important elements and structures in data^[18]^. PCA analysis can extract the two coordinate axes that reflect the difference between samples to the greatest extent, so that the difference of multidimensional data can be reflected on the two-dimensional coordinate graph, and then reveal the simple rules under the complex data background. The more similar the community composition of the samples is, the closer they are in the PCA diagram.

The abscissa in the PCA diagram represents the first principal component, while the percentage represents the contribution value of the first principal component to sample differences. The ordinate represents the second principal component, and the percentage represents the contribution of the second principal component to the sample difference. Each dot in the diagram represents a sample, and samples from the same group are represented in the same color.

It can be concluded from the PCA analysis diagram that the composition of microbial population of each sample in the Experimental group and the Blank group showed obvious differences and differentiation. With the exception of S.Y.4 sample (which may be caused by uneven distribution of soil microbial population), all other samples in the Experimental group showed similarity and dense distribution of loci.

The loci distribution of S.Y.2 and S.Y.3 samples were very concentrated and close to each other, showing a high degree of similarity. The loci distribution of S.Y.1 and S.Y.5 samples were very concentrated and close to each other, showing a high degree of similarity.

##### 2.3.2.3 NMDS analysis

NMDS analysis is also known as Non-Metric Multi-Dimensional Scaling statistics^[19]^, which is a ranking method suitable for ecological studies. NMDS is a nonlinear model based on Bray-Curtis distance. It is designed to overcome the shortcomings of linear models (including PCA and PCoA) and better reflect the nonlinear structure of ecological data^[20]^.

According to the species information contained in the samples, NMDS analysis reflects the species information in the form of points in the multidimensional space. The degree of difference between different samples reflects the inter-group and intra-group differences of samples by the distance between points. Each point in the NMDS analysis chart represents a sample, the distance between points represents the degree of difference, and samples of the same group are represented by the same color. When Stress is less than 0.2, it indicates that NMDS can accurately reflect the degree of difference between samples.

According to the NMDS analysis chart, the difference degree of microbial community composition of each sample in the Experimental group and the Blank group showed obvious differentiation.

The difference between samples in Blank group was significantly greater than that in Experimental group. In the Experimental group, except for S.Y.4 sample (which may be caused by uneven distribution of soil microbial population), the differences among other samples were not large and the distribution of loci was very dense.

The differences among S.Y.1, S.Y.2 and S.Y.3 samples were very small, and the distribution of loci almost overlapped. This indicates that the uniformity and consistency of microbial population composition of the Experimental group is significantly higher than that of the Blank group.

### 2.4 UPGMA clustering tree analysis

In order to study the similarity between different samples, cluster analysis can also be carried out on samples to build a cluster tree of samples. In environmental biology, UPGMA (Unweighted pin-group Method with Arithmetic Mean)^[21]^ is a commonly used clustering analysis Method, which was originally used to solve the classification problem. The basic method of UPGMA is: first gather the two samples with the smallest distance together, and form a new node (new sample), whose branch point is located at 1/2 of the distance between the two samples. Then calculate the average distance between the new “sample” and other samples, and find out the minimum 2 samples for clustering. This is repeated until all the samples are clustered together and a complete cluster tree is formed.

Weighted Unifrac distance matrix and Unweighted Unifrac distance matrix are usually used for UPGMA clustering analysis, and the clustering results are integrated with the relative species abundance of each sample at the gate level.

#### 2.4.1 UPGMA vclustering analysis based on Weighted Unifrac distance

According to the analysis of the inter-group evolutionary tree based on Weighted Unifrac distance (FIG. 10), except for K.B.2 sample in the Blank group and S.Y.4 sample in the Experimental group (possibly due to the uneven distribution of soil microbial population), the microbial populations in the Experimental group and the Blank group showed obvious differentiation. And there are two major subgroups.

**FIG. 5.**
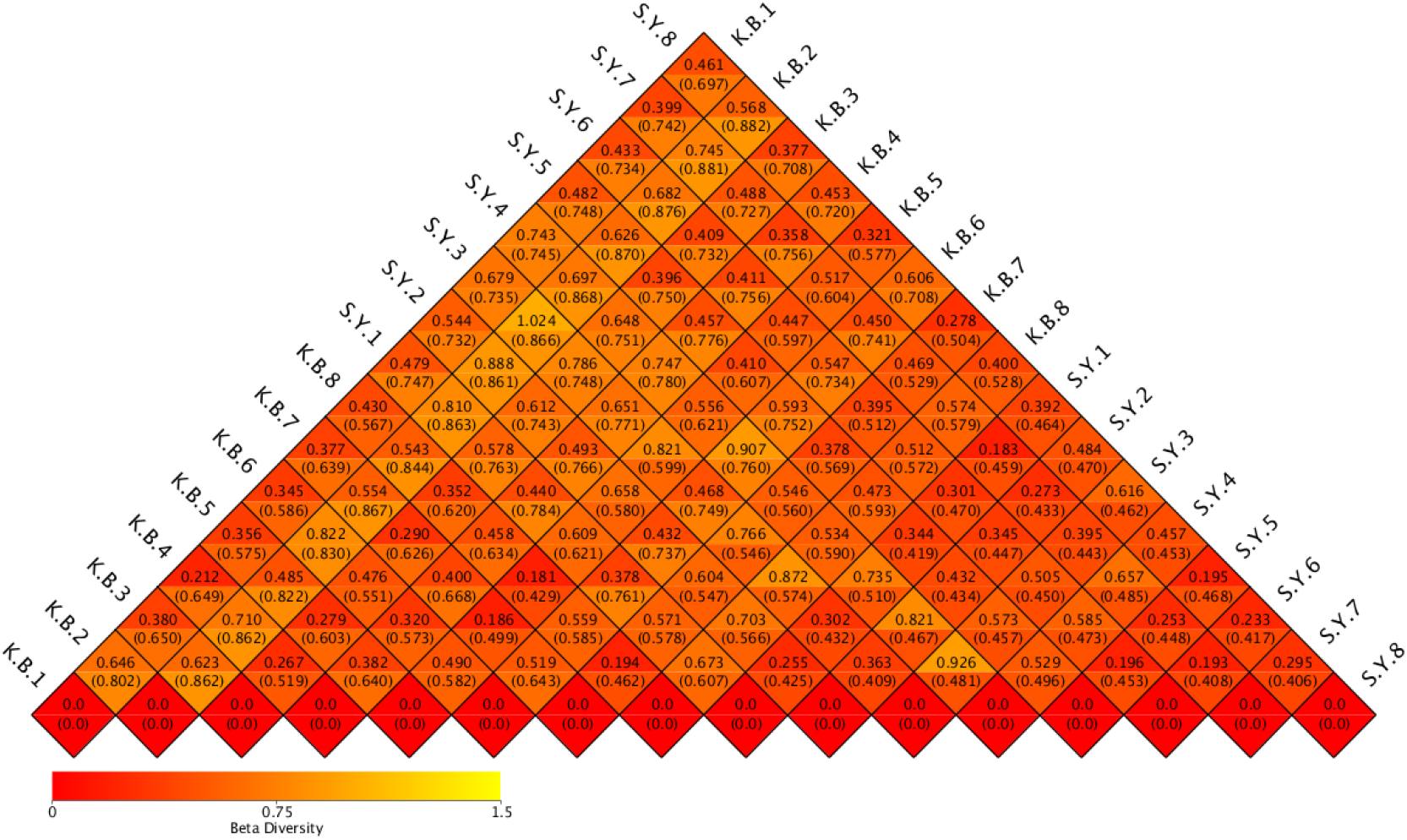
Heatmap of β-Diversity index

**FIG. 6.**
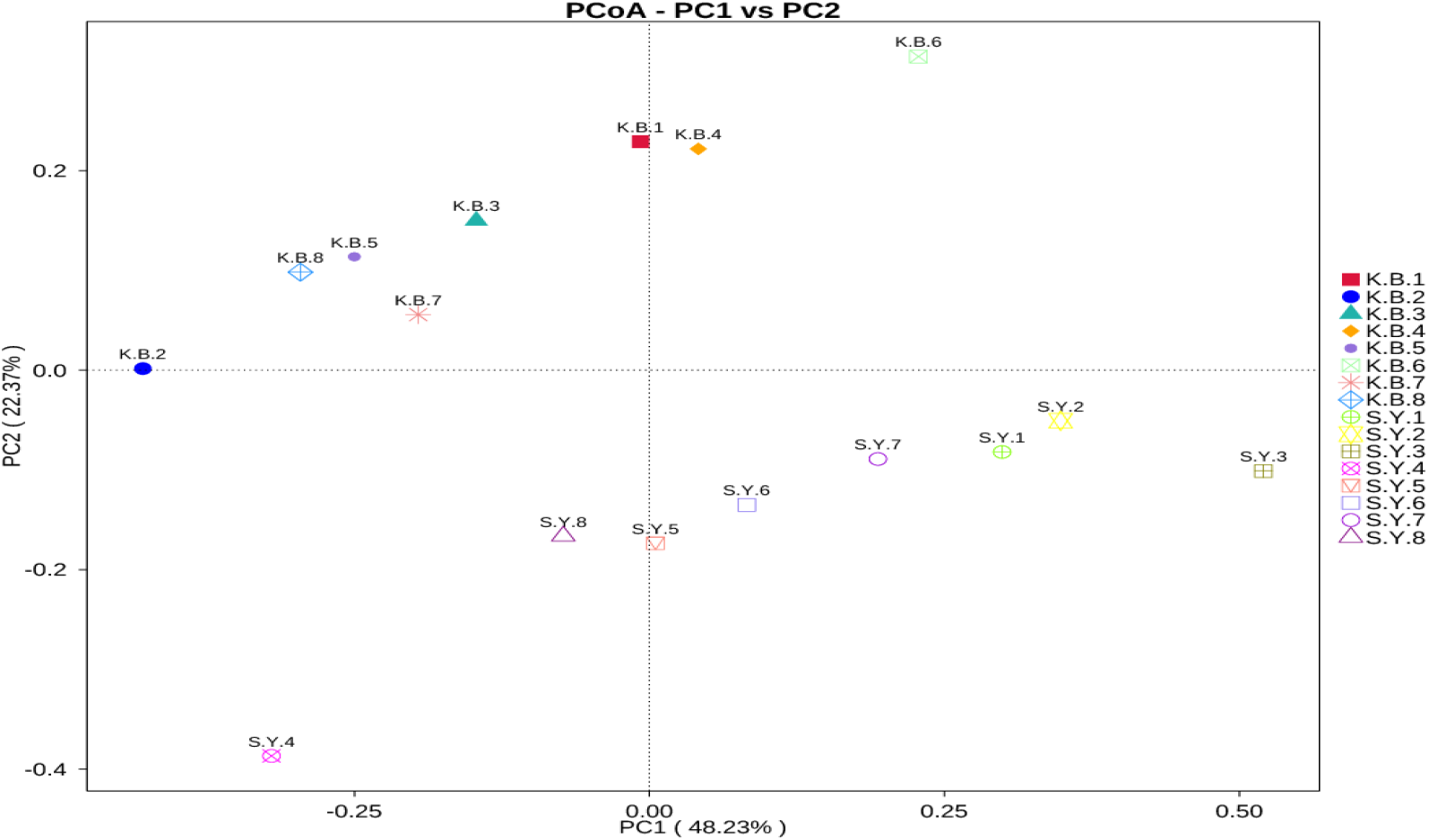
PCoA analysis based on Weighted Unifrac distance

**FIG. 7.**
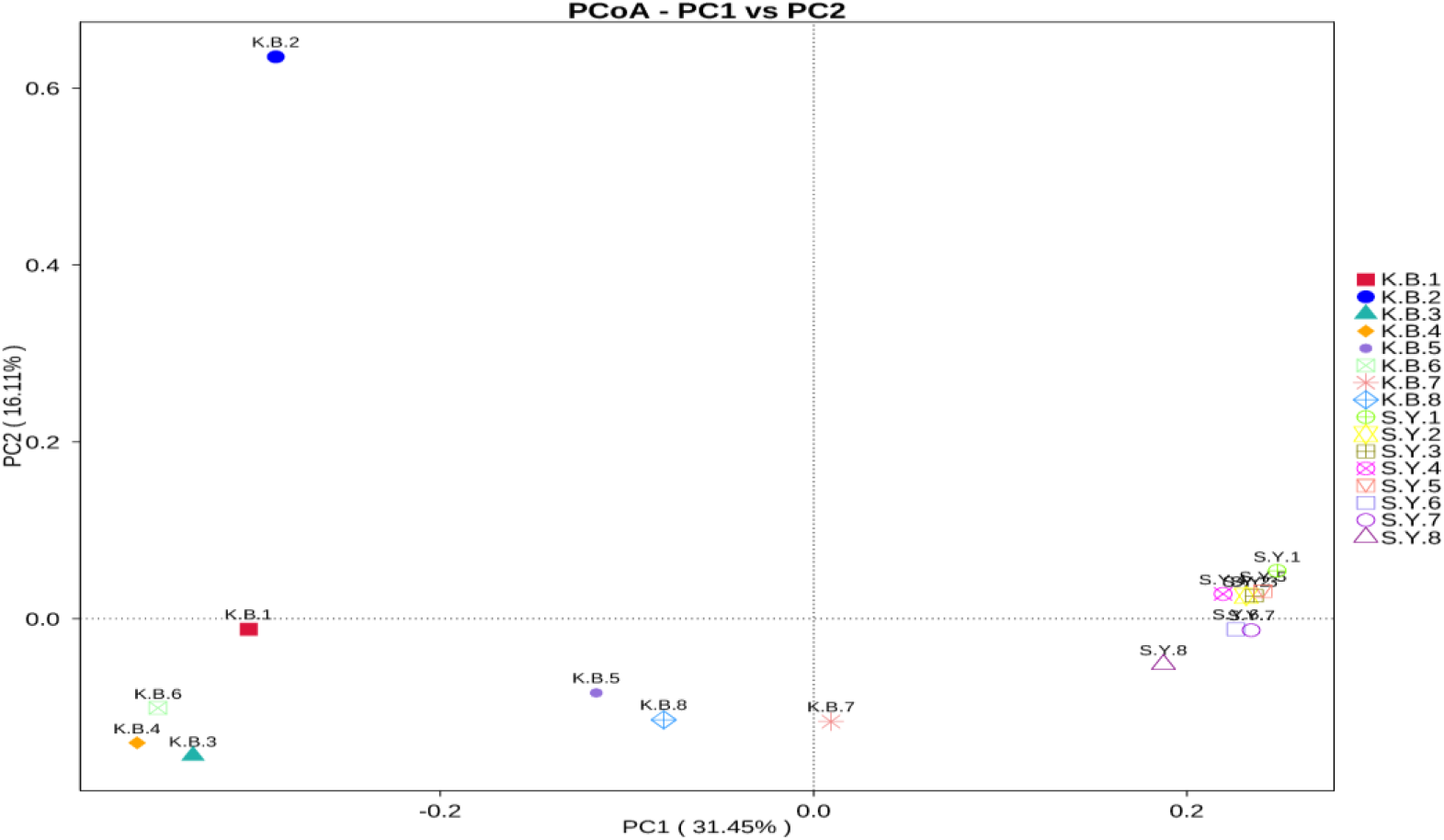
PCoA analysis based on Unweighted Unifrac distance

**FIG. 8.**
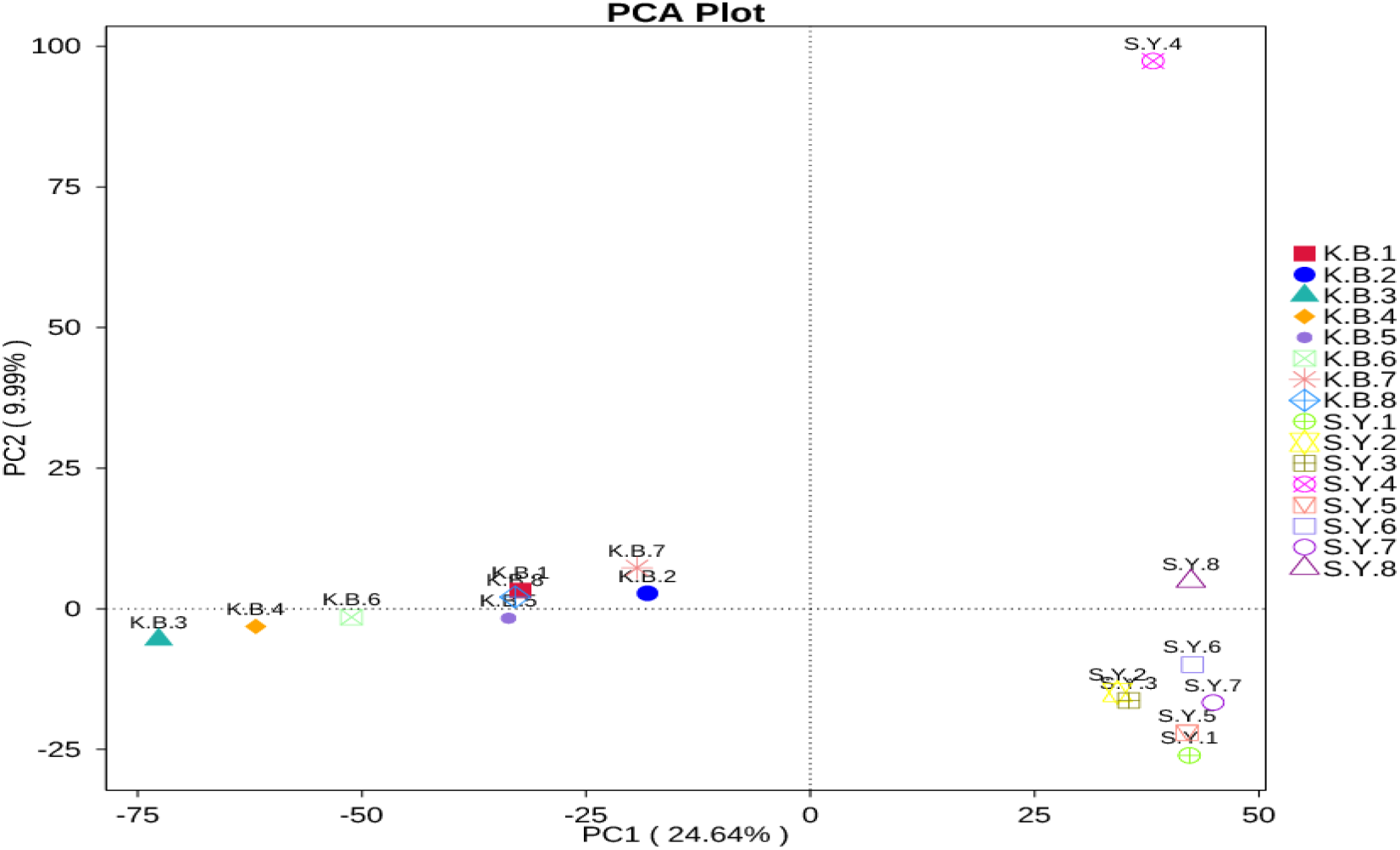
PCA analysis diagram

**FIG. 9.**
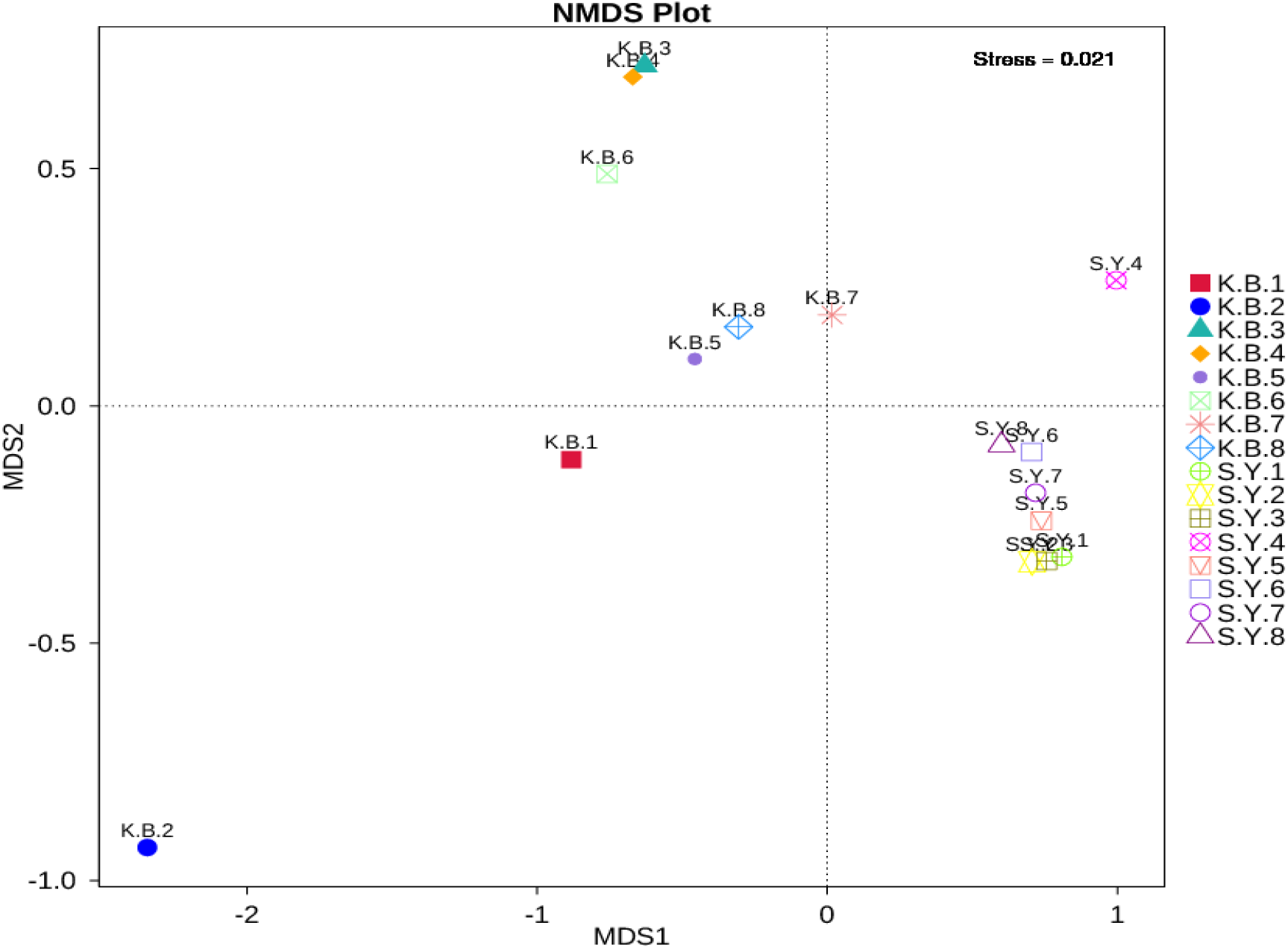
NMDS analysis diagram

**FIG. 10.**
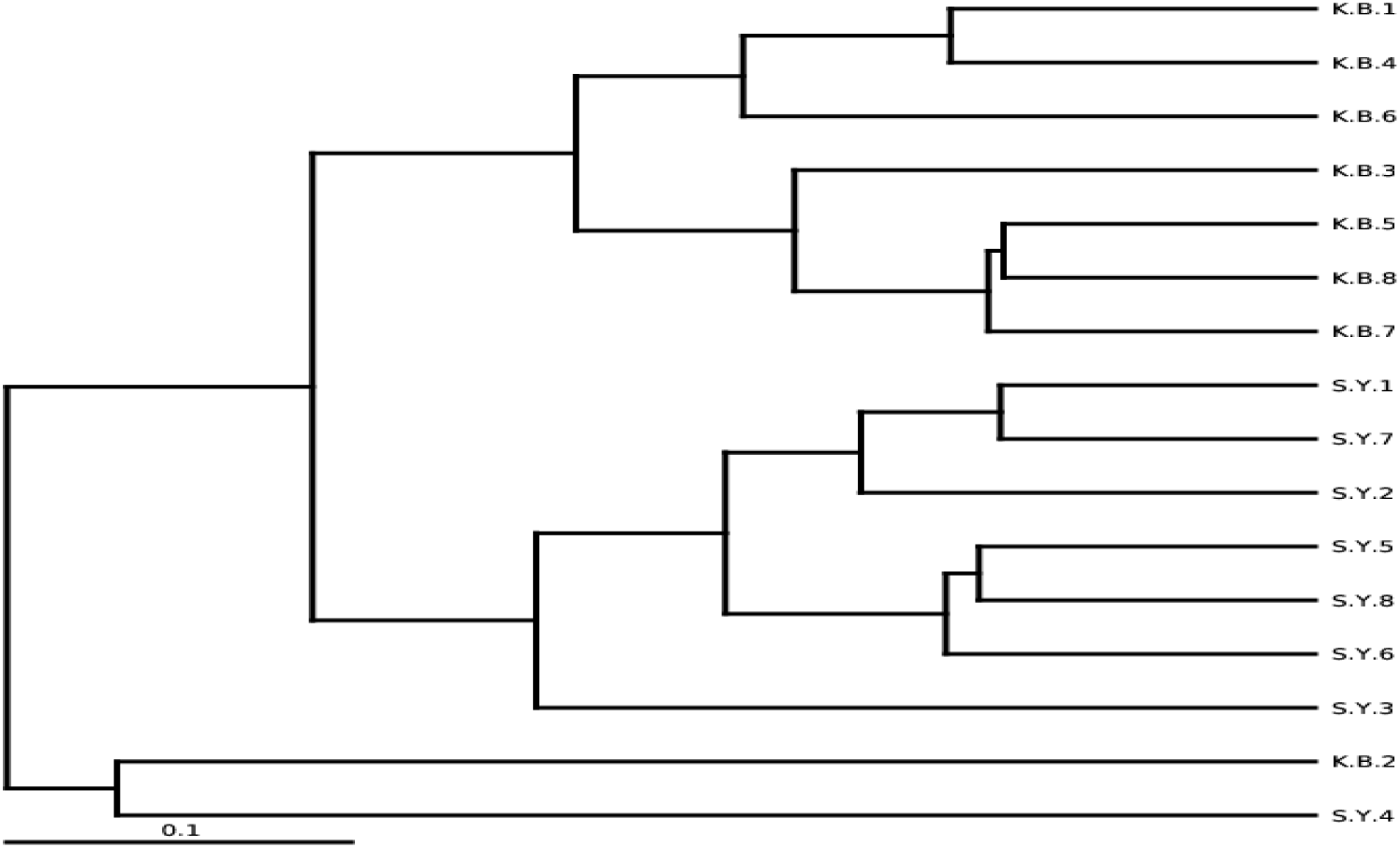
Inter-group evolutionary tree based on Weighted Unifrac distance

K.B.1 and K.B.4, K.B.5 and K.B.8 in the Blank group, and S.Y.1 and S.Y.7, S.Y.5 and S.Y.8 in the Experimental group showed high degree of affinity and homology. On the whole, all samples in the Blank group (except K.B.2 sample) and all samples in the Experimental group (except S.Y.4 sample) showed relatively consistent intergroup affinity and homology.

Based on the UPGMA clustering tree based on Weighted Unifrac distance (FIG. 11), it can be concluded that the composition of microbial population in the Experimental group showed significant fluctuation compared with that in the Blank group. The population abundance of Firmicutes in the Experimental group decreased, but that of Desulfobacterota and Acidobacteriota increased obviously.

**FIG. 11.**
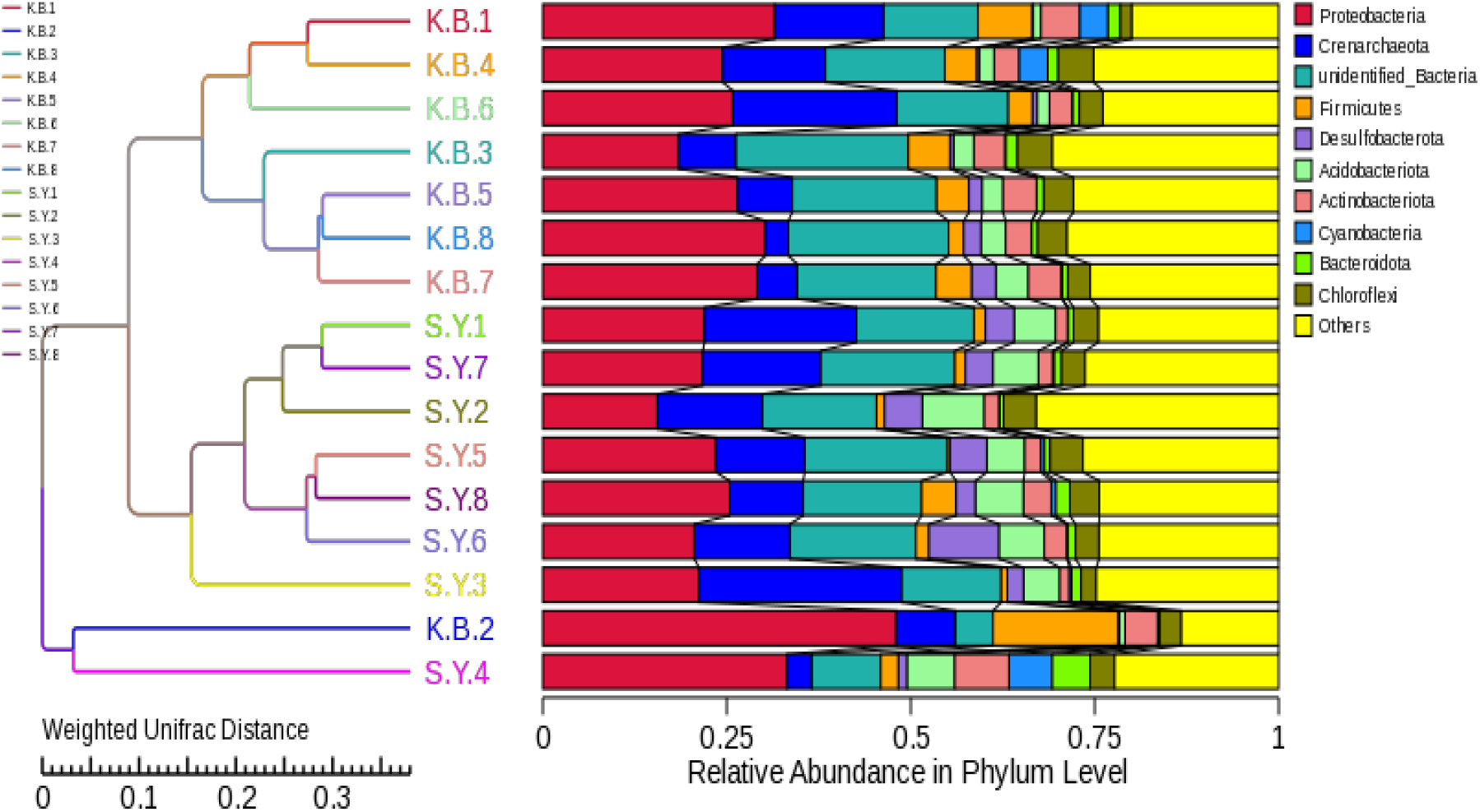
UPGMA clustering tree based on Weighted Unifrac distance

#### 2.4.2 UPGMA clustering analysis based on Unweighted Unifrac distance

According to the analysis of the inter-group evolutionary tree based on Unweighted Unifrac distance (FIG. 11), except for K.B.2 samples in the Blank group (possibly due to uneven distribution of soil microbial population), the microbial populations in the Experimental group and the Blank group showed obvious differentiation. On the basis of UPGMA clustering analysis based on Weighted Unifrac distance, three subgroups are further differentiated.

Among them, K.B.3 and K.B.4, K.B.5 and K.B.8 in the Blank group, S.Y.1 and S.Y.5, S.Y.2 and S.Y.3, S.Y.7 and S.Y.8 in the Experimental group showed high degree of affinity and homology. On the whole, all samples in the Blank group (except K.B.2 sample) and all samples in the Experimental group showed relatively consistent inter-group affinity and homology.

According to the UPGMA clustering tree based on Unweighted Unifrac distance (FIG. 13), the microbial population composition of the Experimental group showed significant fluctuations compared with that of the Blank group, and the population abundance of Firmicutes in the Experimental group decreased. The abundance of Acidobacteriota increased significantly.

**FIG. 12.**
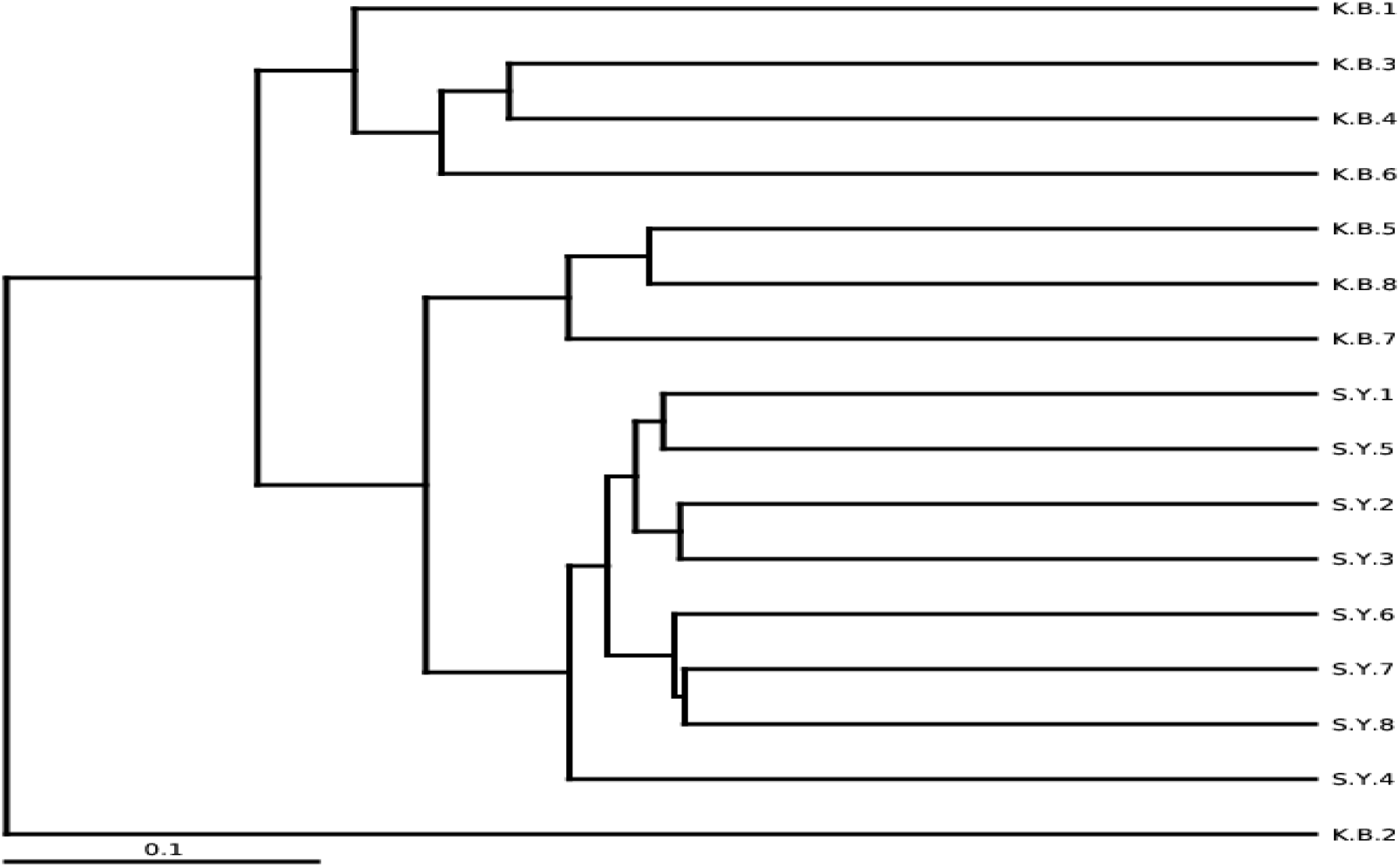
Inter-group evolutionary tree based on Unweighted Unifrac distance

**FIG. 13.**
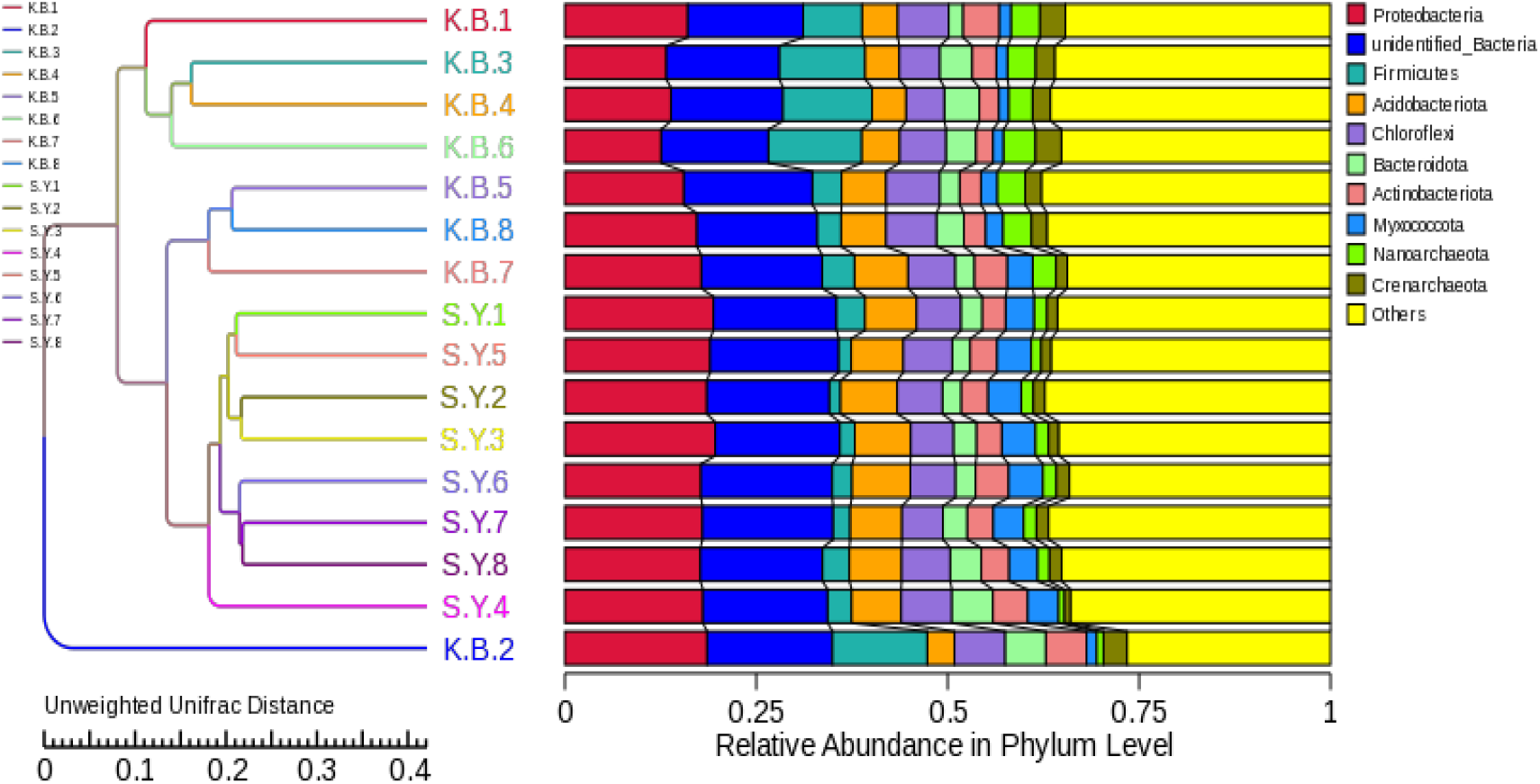
UPGMA clustering tree based on Unweighted Unifrac distance

## 3 Discuss

Through the above analysis, it can be concluded that the abundance of microbial population of soil samples in the Experimental group planted with JunCao “Oasis NO. 1” is significantly higher than that in the blank section without JunCao “Oasis No. 1”. These results indicated that the JunCao “Oasis NO. 1” could effectively improve the microbial population abundance of saline-alkali soil and restore the microbial population ecosystem of saline-alkali soil. Therefore, it has good characteristics of soil improvement and management in saline-alkali land.

In terms of the composition of soil microbial population in the Experimental group, the microbial population in the Experimental group planted with JunCao “Oasis NO. 1” was compared with that in the blank lot without JunCao “Oasis No. 1”, not only the abundance of microbial population increased, More importantly, the composition of microbial community in the Experimental group planted with JunCao “Oasis NO. 1” also showed obvious differentiation.

Among the phylogenetic trees of each genus, Pseudomonas from Proteobacteria was the dominant genus in the soil samples of the Experimental group, and there was a large number of Pseudomonas in the soil samples of the Experimental group. However, the dominant position of Sphingomonas was not obvious, and only S.Y.4 sample from the Experimental group had a large distribution in Sphingomonas.

In the Weighted Unifrac distance intergroup evolutionary tree, the microbial populations of Experimental and blank soil samples were divided into two groups. In the inter-group evolutionary tree based on Unweighted Unifrac distance, the soil samples in the Experimental group and the blank group differentiated into three major groups, while the soil samples in the Experimental group further differentiated into two major groups.

The microbial population composition of soil samples in the Experimental group showed significant changes compared with that in the Blank group. The population abundance of Firmicutes in the Experimental group decreased, but that of Desulfobacterota and Acidobacteriota increased obviously. Acidobacteria is a new bacterium established in recent years and has important ecological effects. This fully indicates that JunCao “Oasis 1” can not only effectively improve the abundance of microbial population in salt-alkali soil, but more importantly, it can effectively improve the abundance of beneficial bacteria with important ecological functions in soil. In the soil improvement and management of saline-alkali land, JunCao “Oasis No. 1” has important ecological application value.

Governance alkali-saline land areas in the future, bioremediation method will get more and more widely used, in this paper, by using 16SrDNA high-throughput sequencing methods to analyze saline-alkali soil planting JunCao “Oasis No. 1”changes before and after the differences of soil microbial diversity of species abundance, proved JunCao “Oasis No. 1” has excellent characteristics of saline soil, It provides a new method and idea for treating saline-alkali soil by biological method, and has a good practical guiding significance and application value.

